# Identification and characterisation of splicing regulators in *Toxoplasma gondii*

**DOI:** 10.1101/2021.06.27.450092

**Authors:** V Vern Lee, Simona Seizova, Paul J. McMillan, Emma McHugh, Christopher J. Tonkin, Stuart A. Ralph

## Abstract

The splicing of mRNA constitutes a major source of co- and post-transcriptional regulation in metazoans. In particular, members of the serine/arginine (SR) protein family are essential splicing factors that are implicated in the regulation of gene expression and RNA metabolism. However, very little is known about these proteins in apicomplexans, a phylum that includes some of the most important global parasites. In this study, we investigated the suite of three uncharacterised SR proteins in *Toxoplasma gondii* and show that all three are found localised to nuclear speckles. We show, by genetic ablation, that *Tg*SR1 is particularly important for *T. gondii* growth. Using RNA-seq, we also characterised the global gene expression and splicing regulation of these proteins. We find that the SR proteins regulate several types of alternative splicing of distinct but overlapping subsets of transcripts, as well as impacting transcript abundance. Most of the alternative splicing events are non-productive intron retention events that do not appear to affect transcript abundance. The splicing sites of the impacted transcripts are enriched in characteristic SR binding motifs. We also identified and conditionally knocked down two putative kinases of SR proteins. The kinases are localised to nuclear speckles and are essential to parasite survival. Their perturbation resulted in widespread changes to splicing, but the affected transcripts did not mirror the patterns seen in knockouts of individual SRs, suggesting an absence of a simple relationship between SRs and these putative kinase regulators. Overall, this study reveals a complex system of splicing factors and kinases that post-transcriptionally regulate gene expression in *T. gondii*.

## Introduction

The phylum Apicomplexa is a group of intracellular parasitic protists that includes some of the most important parasites impacting human and veterinary health, such as *Plasmodium and Toxoplasma*. *Plasmodium falciparum,* the pathogen responsible for most severe malaria, infects over 200 million individuals and kills 400,000 each year (1). Similarly, *Toxoplasma gondii*, the causative agent of toxoplasmosis, is a widespread disease, with over a third of the world population estimated to be infected, and seropositivity rates of up to 90% in some countries (2). Toxoplasmosis remains a significant threat to immunocompromised, young, or pregnant individuals, who are at risk of developing severe pathology (3, 4).

The lifecycle of most apicomplexans is complex, and the parasites must contend with dynamically changing environments and hosts. Central to the parasites’ success is the tight and constant reprogramming of gene expression underlying parasite development. Studies on *P. falciparum* have revealed developmentally distinct transcriptomic and proteomic patterns as the parasite transitions between life cycle stages. (5, 6). Similarly, the transition between *T. gondii* acute-stage tachyzoites and chronic, cyst-forming bradyzoites requires significant changes to transcript abundance for most genes (7). Such changes require an extensive network of regulatory mechanisms, such as transcriptional, post-transcriptional, and epigenetic control of genes (8).

RNA splicing plays a major role in transcriptional and post-transcriptional control in metazoans, but the process is much less understood in apicomplexan parasites, and indeed in any protist species. Multi-exon genes require the constitutive removal of introns from pre-mRNA and the ligation of exons to produce mature mRNA transcripts. However, alternative splicing (AS) can occur, where a deviation in splicing patterns results in multiple transcript isoforms. This allows a single gene to encode different protein isoforms with different molecular and cellular functions, but in many cases can also generate non-functional products. Indeed, many genes have been shown to produce multiple protein isoforms with altered structure, activity, modification and localisation (9, 10). AS can also directly impact gene expression, for example by altering small-RNA binding sites (11) or through the nonsense-mediated RNA decay (NMD) pathway for non-productive transcripts (12). The NMD pathway in particular regulates up to 25% of transcripts in humans, some a result of splicing mistakes, and some a result of targeted regulation (12).

A key element in the control of alternative splicing in metazoans is the serine/arginine-rich (SR) protein family. SR proteins are defined by the presence of extended arginine and serine (RS) dipeptides regions and at least one RNA recognition motif (RRM) (13). SR proteins mediate AS by binding to specific regions of pre-mRNA, and enhancing or inhibiting the interactions between components of the spliceosome and proximal splice sites (14). Thus, SR proteins can enhance or repress constitutive/alternative splicing depending on the context (15). RNA binding sites of SR proteins are often poorly defined. The binding sequences have been established to be purine rich but are degenerate, at least at a primary sequence level, due in part to the seemingly accommodating structure of the RRM domains (14). For example, the human SR protein SRSF2 has a novel structure that allows equal recognition of guanines and cytosines (16). The poor definition allows the SR proteins to generate a broad SR-RNA interactome, which is necessary for the efficient splicing of divergent protein-coding sequences (17). Moreover, the RS domains of SR proteins often interact with other RS domain-containing proteins (18). Therefore, cooperation or competition between SR proteins is vital in determining the final spliced transcripts. While primarily known to be modulators of AS, SR proteins have also been implicated in other roles such as regulating genomic stability, transcription, mRNA transport, and translation (19).

In line with their function, SR proteins typically localise to the nucleus in subnuclear regions called speckles. Speckles are subcellular domains for the storage and assembly of splicing factors (20). A speckled-like localisation is strongly indicative of proteins involved in RNA splicing; virtually all splicing factors are known to localise to nuclear speckles (21). Nuclear speckles organise transcriptionally active genes at their periphery, where splicing factors are shuttled to for their action (22). The localisation and specific activity of SR proteins require phosphorylation of the RS domain (23). This process is facilitated by multiple kinases from the CMGC superfamily of kinases, particularly, from the SRPK, CLK, and DYRK subfamily (24). In humans, 3 SRPKs (SRPK1–3), 4 CLKs (CLK1–4), and DRYK1a have been shown to be involved in SR protein phosphorylation mainly through genetic ablation studies (24). An additional CLK kinase, PRP4, appears to phosphorylate SR proteins in *Schizosaccharomyces pombe* (25) as well, though the role of PRP4 in metazoans is less clear (25). SR proteins and their regulators have been revealed to be essential in varied life stages of different *Plasmodium* species, and several inhibitors have been validated (26–29), but very little is known about these proteins in *T. gondii*. Four putative SR proteins were previously identified but only one (*Tg*SR3) has been characterised (30). Even less is known about the putative kinases that modulate the process. Putative *T. gondii* SRPK, CLK and DYRK genes have been identified through in silico analyses (31), but have not been characterised.

In this study, we attempted to broadly characterise all putative *T. gondii* SR proteins and kinases of SR proteins. We demonstrate that all three previously uncharacterised putative SR proteins localise to nuclear speckles, and that genetic ablation of the proteins results in changes to gene expression and AS, as identified by RNA-seq analysis. This global approach revealed that different SR proteins regulate subsets of AS to different extents, but affected the same types of AS events (i.e., intron retention, alternative 5’or 3’ splice site change, and exon skipping) in similar proportions. Only ablation of one of the SR proteins, *Tg*SR1, results in a detectable fitness cost in *in vitro* growth, which we attribute to a general splicing defect. Differential AS events were enriched in sequence motifs that are characteristic of AS modulator-binding sequences, further indicating their role as AS factors. We also identified five putative kinases of SR proteins, and successfully adapted the auxin-inducible degron system to localise and conditionally knock down three of them. Two of these, which are homologues of human CLK and PRP4 kinases respectively, localise to nuclear speckles and are essential to parasite survival. We further characterise the two using RNA-seq analyses and find that they extensively regulate gene expression and splicing events. Unusually, the transcriptomic changes that arise from these ablations overlapped poorly with that of the SR proteins characterised in this study. The mismatch indicates the complex nature of the splicing regulation in *T. gondii* required for completion of a multifaceted life cycle.

## Results

### *Tg*SRs localise to nuclear speckle-like structures

We had previously identified 4 putative *T. gondii* SR homologues (*Tg*SR1-4) based on the RRM sequences of the 12 known human SR proteins, and characterised one of them, *Tg*SR3 (30). However, as there is no clear direct orthology between members of the family, we sought to characterise and determine if the other putative *Tg*SR proteins (*Tg*SR1-TGME49_319530, *Tg*SR2-TGME49_217540 and *Tg*SR4-TGME49_270640) are alternative splicing factors. To investigate the localisation of these proteins, we genetically inserted three HA tags and drug selection cassette at the 3’end of the endogenous genes using an established CRISPR/Cas9 approach (32, 33) in PruΔ*ku80* parasite strain (Fig. 1A). Selected clonal transgenic parasites were genotyped by PCR using primers specific to the selectable marker and 3’ UTR of the individual genes to verify insertion (Fig. 1A-B). Western blotting analyses confirmed the expression of each of the tagged proteins, which corresponded to their expected molecular weight (Fig. 1C). SR proteins are typically found localised to nuclear speckles (21). Widefield immunofluorescence assays (IFAs) with deconvolution for each of the HA-tagged putative *T. gondii* SR proteins revealed a speckle-like signal which overlaps the nuclear DAPI signal, with apparent absence from the nucleolus (which has a weak DAPI signal) (Fig. 1D). It was unclear from these images if the speckled pattern might be reticulated, which would be unusual for splicing factors. Thus, to obtain increased resolution, we utilised 3D Structured Illumination Microscopy (3D-SIM). The 3D-SIM images confirmed a strong nuclear speckle-like localisation with no further microstructure (Fig. 1E). Interestingly, the nuclear speckles formed a bifurcated pattern in some of the parasites (Figure 1E right), which resembles the nucleus during endogeny/mitosis. This differs from human nuclear speckles that become dispersed and form mitotic interchromatin granules (MIGs) in the cytoplasm during mitosis (34). Nevertheless, the localisation of the proteins within nuclear speckles supports their involvement in the splicing process.

**Figure 1.**
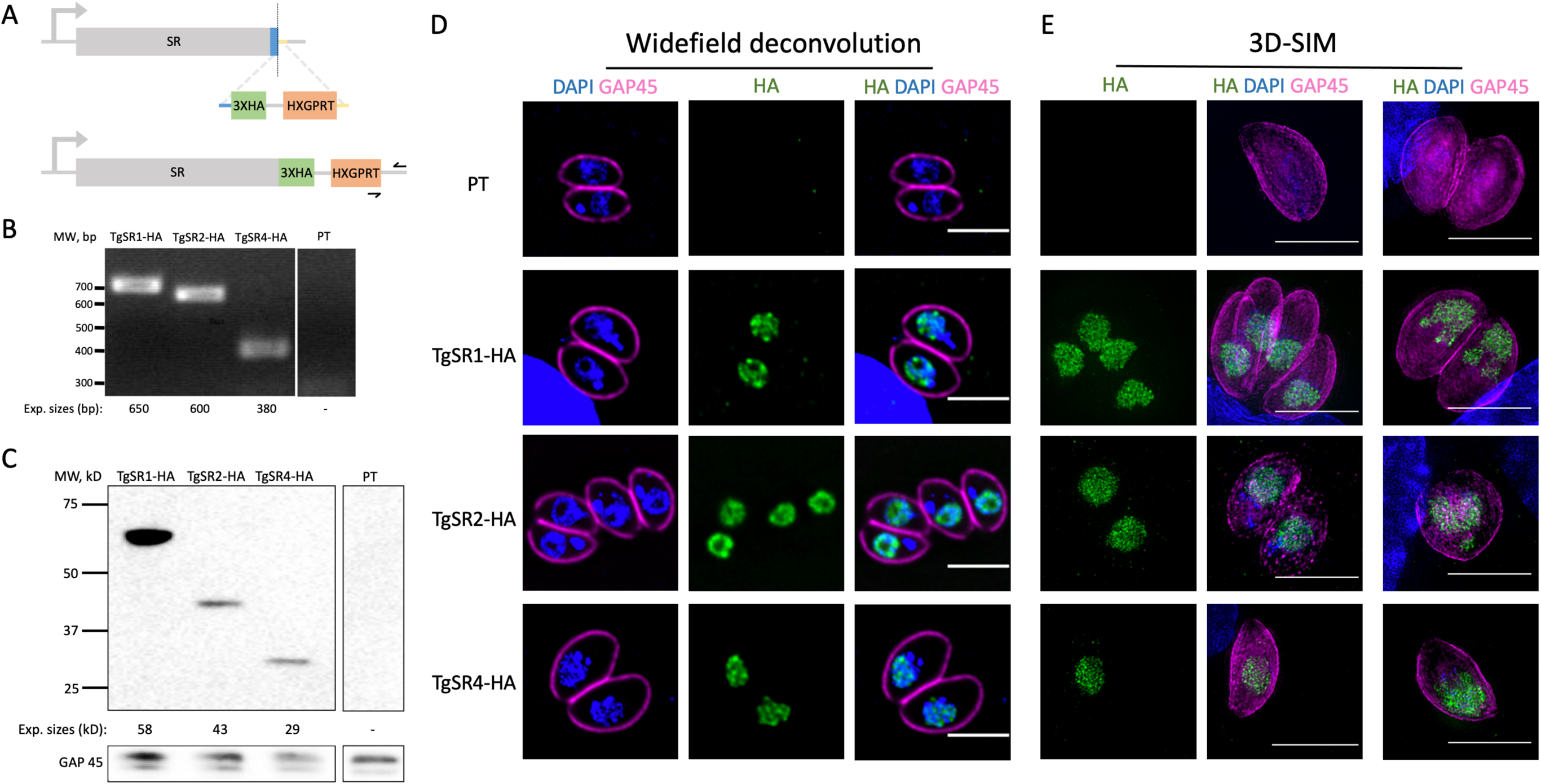
**Localisation of *Tg*SR proteins.** (A) Schematic for the generation of *Tg*SR-HA tagged parasite lines. HXGRPT is the selection marker, blue and yellow segments represent homology arms for homology-directed repair of the Cas9-induced double strand break (vertical dotted line). Black arrows represent primer targets for PCR genotyping (B) PCR genotyping of *Tg*SR-HA parasites. (C) Western blot of total protein purified from *Tg*SR-HA parasites probed with anti-HA and anti-GAP45 antibodies. (D-E) Representative immunofluorescence microscopy images of *Tg*SR-HA intracellular parasites stained with DAPI (blue), anti-HA (green) and anti-GAP45 (magenta) antibodies. Images are maximum projections of wide-field deconvoluted (D) or 3D-SIM (E) z-stacks. Scale bars = 3 µm. The parental line (PT) was used as the negative control in each (B-E) assay.

### TgSRs are not essential in tachyzoite and bradyzoite stages of T. gondii

Based on a previous genome wide CRISPR screen (35), the three *Tg*SR proteins are predicted to be dispensable for parasite *in vitro* growth. We therefore attempted to directly knock out the genes by simply inserting a drug selectable cassette (DHFR) (36) within the first exon in the HA-tagged mutants. However, only the expression of *Tg*SR2-HA appeared to be completely ablated despite repeated attempts. Because of this refractoriness to ablation, we resorted to homologous replacement of the near full-length *Tg*SR1/4-HA genes with the DHFR cassette (Fig. 2A). PCR genotyping of selected monoclonal transgenics confirmed that the DHFR cassettes were correctly inserted in each case (Fig. 2B). We then performed western blotting analyses and IFAs on the mutants and found that the previously intact HA signals were no longer detectable (Fig. 2C-D), indicating that the genes were indeed disrupted and no longer expressed. Successfully generating the mutants validated the prediction that the genes were not essential to parasite survival at the tachyzoite stage.

**Figure 2.**
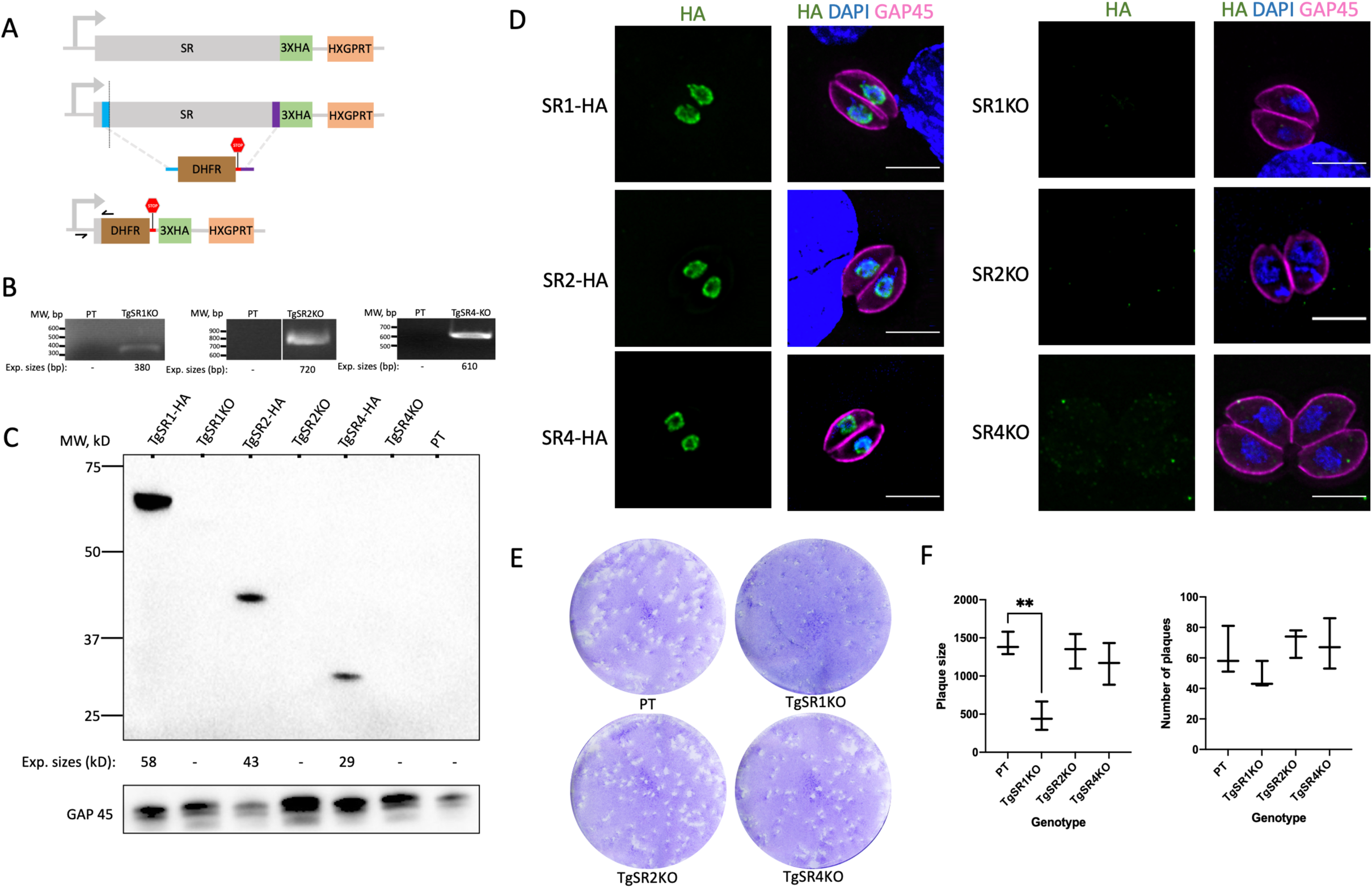
**Ablation of *Tg*SR proteins.** (A) Schematic for the generation of *Tg*SR-HA-knockout (KO) parasite lines. HXGRPT and DHFR are selection markers. Black arrows represent primer positions for PCR genotyping. (B) PCR genotyping of *Tg*SR-HA-KO parasites. (C) Western blot of total protein purified from *Tg*SR-HA and *Tg*SR-HA-KO parasites probed with anti-HA and anti-GAP45 antibodies. (D) Representative immunofluorescence microscopy images of *Tg*SR-HA and *Tg*SR-HA-KO intracellular parasites stained with DAPI (blue), anti-HA (green) and anti-GAP45 (magenta) antibodies. Images are maximum projections of wide-field deconvoluted z-stacks. Scale bars = 3 µm. (E) Representative plaque assay images of *Tg*SR-HA-KO parasites in HFF cell monolayers stained with crystal violet. (F) Quantification and statistical analysis of the mean size (left) and number (right) of plaques from (E) in 3 biological replicates using the Student’s paired t-test. **, P < 0.01. The parental line was used as the negative control in each (B-F) assay.

To determine if knocking out the genes resulted in a growth defect, we performed plaque assays on fibroblast monolayers. As shown in Figure 2E, the assays revealed impaired growth for *Tg*SR1KO but not *Tg*SR2KO and *Tg*SR4KO. Plaque sizes were statistically significantly reduced (p ≤ 0.05) compared to the parental line, but not plaque number (Fig. 2F), which indicates that initial invasion is unimpaired, but that growth rate or number is reduced. No other defects for any of the mutants were detected at this stage. We had previously shown that AS is stage-specific in *P. berghei* and that one of the *Pb*SR proteins is required for the differentiation of male gametes (26). Changes in AS also occur when *T. gondii* tachyzoites differentiate into bradyzoites (7) and so we tested whether the *Tg*SR proteins were required for differentiation into bradyzoites. We differentiated our mutants using alkaline stress and performed IFAs 7 days post induction. All the mutants were able to differentiate into bradyzoites as indicated by the detection of bradyzoite specific markers with no observable defects compared to the parental line (Fig. S1).

### Phylogenetic analysis identifies *T. gondii* homologues of kinases that phosphorylate SR proteins

In metazoans, multiple related kinases are required to phosphorylate SR proteins for their localisation and function. However, very little is known about these genes in *T. gondii.* Using the protein sequence of the nine human SR proteins kinases (SRPK1–3, CLK1–4, PRP4 and DRYK1a), we retrieved and aligned homologues from *S. pombe*, *Arabidopsis thaliana*, *T. gondii* and *P. falciparum*, and constructed a phylogenetic tree from this alignment (Fig. 3A). Interestingly, while the human and *A. thaliana* SRPK and CLK families each consists of multiple paralogs, only one SRPK and one CLK copy could be identified in *T. gondii* and *P. falciparum*, as has been previously found (31). This is similar to *S. pombe* in which only a single SRPK or CLK homologue has been identified (37). The SRPK and CLK homologues all resolved within their respective family clade. Single *T. gondii* PRP4 and DYRK1a homologues could also be identified based on clustering within the same clade of the other PRP4/DYRK1a homologues. We refer to the *T. gondii* genes identified in this analysis using the name of whichever clade family they resolved in (*Tg*SRPK, *Tg*CLK, *Tg*PRP4, *Tg*DYRK1a). Unexpectedly, we find an unresolved clade consisting of one gene each from *T. gondii* (TGME49_204280) and *P. falciparum* (PF3D7_1443000). The *P. falciparum* gene was annotated in PlasmoDB (38) as SRPK2 and had been previously referred to in a publication as *Pf*CLK2 (28). On the other hand, the corresponding *T. gondii* orthologue was annotated as a putative DYRK. Talevich and colleagues (31) had previously suggested that the *Plasmodium* gene is atypical in that it has characteristics of both SRPK and DYRK families. To avoid confusion, we refer to the *T. gondii* gene as *Tg*DYRK as it appears most closely related to the DYRK1a clade.

**Figure 3.**
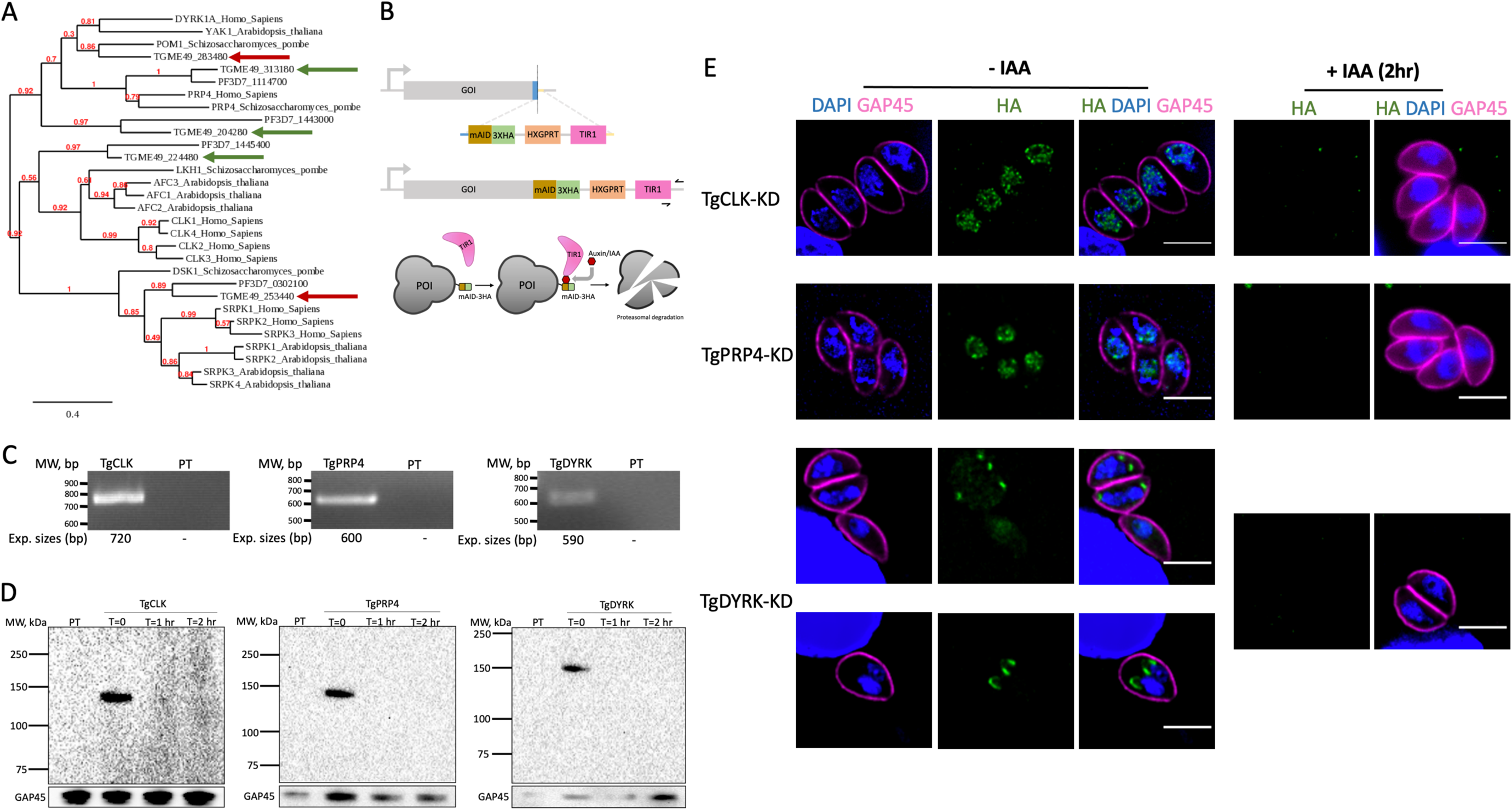
**Phylogenetic analysis and AID-HA tagging of putative SR kinases in *T. gondii*.** (A) Phylogenetic tree for putative kinases of SR proteins in *H. sapiens*, *S. pombe*, *A. thaliana*, *gondii* and *P. falciparum*. Numbers in red are branch support values derived with the approximate likelihood ratio test (aLRT). Scale bar of branch length is proportional to the number of substitutions per site. Arrows represent *T. gondii* genes that were subsequently successfully (green) or unsuccessfully (red) AID-HA tagged. (B) Schematic of the strategy used to adapt the auxin inducible degron system for the putative kinases of SR proteins in *T. gondii*. HXGRPT is the selection marker. Black arrows represent primer targets for PCR genotyping. (C) PCR genotyping of AID-HA tagged parasites. (D) Western blot of total protein purified from AID-HA tagged parasites before and after the addition of IAA at fixed timepoints, probed with anti-HA and anti-GAP45 antibodies. (E) Representative immunofluorescence microscopy images of AID-HA tagged intracellular parasites in the absence or presence of IAA, stained with DAPI (blue), anti-HA (green) and anti-GAP45 (magenta) antibodies. Images are maximum projections of wide-field deconvoluted z-stacks. Scale bars = 3 µm. The parental line was used as the negative control in each (C-E) assay.

### *Tg*CLK and *Tg*PRP4 but not *Tg*DYRK localise to nuclear speckle-like structures

With the exception of DYRK1a, all the other *T. gondii* kinases were predicted to be essential to parasite survival (35). Thus, we attempted to utilise the auxin-inducible degron (AID) system to conditionally knock down the proteins. The system requires two transgenic elements-a plant auxin receptor, transport inhibitor response 1 (TIR1), and an AID or mini-AID (mAID) genetically fused to the protein of interest. Addition of auxin/IAA targets the tagged protein for rapid proteasomal degradation in a TIR1 expressing cell line (39). Here, we wanted to combine both elements within a single construct so that only a single transfection and insertion event were needed. We created a plasmid that contains the sequence of mAID fused to 3xHA tags, a downstream selectable marker HXGPRT (hypoxanthine-xanthine-guanine phosphoribosyltransferase), and TIR1, and used the CRISPR/Cas9 approach as before (Fig. 3B). We carried out transfections for each of the putative kinases, but only successfully recovered positive clones as confirmed by PCR genotyping (Fig. 3C) for *Tg*CLK, *Tg*PRP4 and *Tg*DYRK. Western blotting analyses revealed protein bands of expected mass for *Tg*PRP4 (117 kDa) and *Tg*DYRK (140 kDa). However, the observed mass of *Tg*CLK (~120 kDa) was lower than the expected mass of 200 kDa, suggesting that the gene product might be processed post-translationally. In all three clones, addition of IAA resulted in the rapid knockdown of the proteins as indicated by loss of HA signal within 1 hour. This was also observed with IFAs (Fig. 3E). The IFAs revealed the localisation of *Tg*CLK and *Tg*PRP4 to be that of nuclear speckles, again suggesting that they have a role in the splicing process. The localisation of *Tg*DYRK however, was less certain. In some parasites, *Tg*DYRK appeared to have a low intensity diffuse nuclear and cytoplasmic signal. In contrast, in parasites that were undergoing endogeny as indicated by a bifurcated nucleus, a high intensity non-overlapping signal was observed at the peripheral apical end of the bifurcated nucleus. The signal either appeared punctate, or elongated and curved, mimicking the shape of the centrosome. This is similar to the fission yeast DYRK-family kinase POM1 which localises to the cell tips during cell division (40). POM1 is not involved in splicing, but rather in cell morphology, bipolar growth and cytokinesis (41). Due to this and the lack of evidence of DYRK kinases apart from DYRK1a in humans to be involved in the splicing process, we chose to prioritise the other two putative kinases for further analysis in this study.

### *Tg*CLK and *Tg*PRP4 are essential to parasite division

As before, we performed plaque assays on *Tg*CLK and *Tg*PRP4, with or without addition of IAA, to test if the genes were required for parasite survival and growth. In the knockdown condition, tachyzoites from both lines failed to produce any visible plaques (Fig. 4A-B). It has been previously noted that auxin or IAA is non-toxic to *T. gondii* (39). Indeed, addition of IAA to parental (PT) line parasites did not significantly alter parasite plaque size or number compared to the vehicle control (EtOH). The AID-tagged lines with the vehicle control also showed no significant growth difference compared to the parental line. To determine how knocking down the proteins might be impacting parasite survival, we tracked the parasites via IFAs for 2 days. Parasites were allowed to invade HFF cells on day 0 in the presence or absence of IAA. In both conditions, parasites were detected within HFF cells at day 1, indicating that cell invasion could still occur (Fig. 4C). However, there were virtually no dividing/daughter cells present in the IAA induced parasites for both tagged lines compared to the uninduced and parental lines, suggesting a defect in cell division. On day 2, the parental line had continued multiplying and rosettes could be observed in the enlarging parasitophorous vacuoles. On the other hand, the knockdown parasite lines remained as non-divided singlet cells. Moreover, the *Tg*CLK knockdown parasites had lost their characteristic crescent shape and had become rounded, while *Tg*PRP4 knockdown parasites were smaller in size than the controls. Collectively, these results illustrate that these two genes are essential to parasite division and survival at the tachyzoite stage. We attempted to differentiate the parasites into bradyzoites as well following knockdown but failed to observe any by IFA, suggesting that the parasites could not escape the severe phenotype by differentiating.

**Figure 4.**
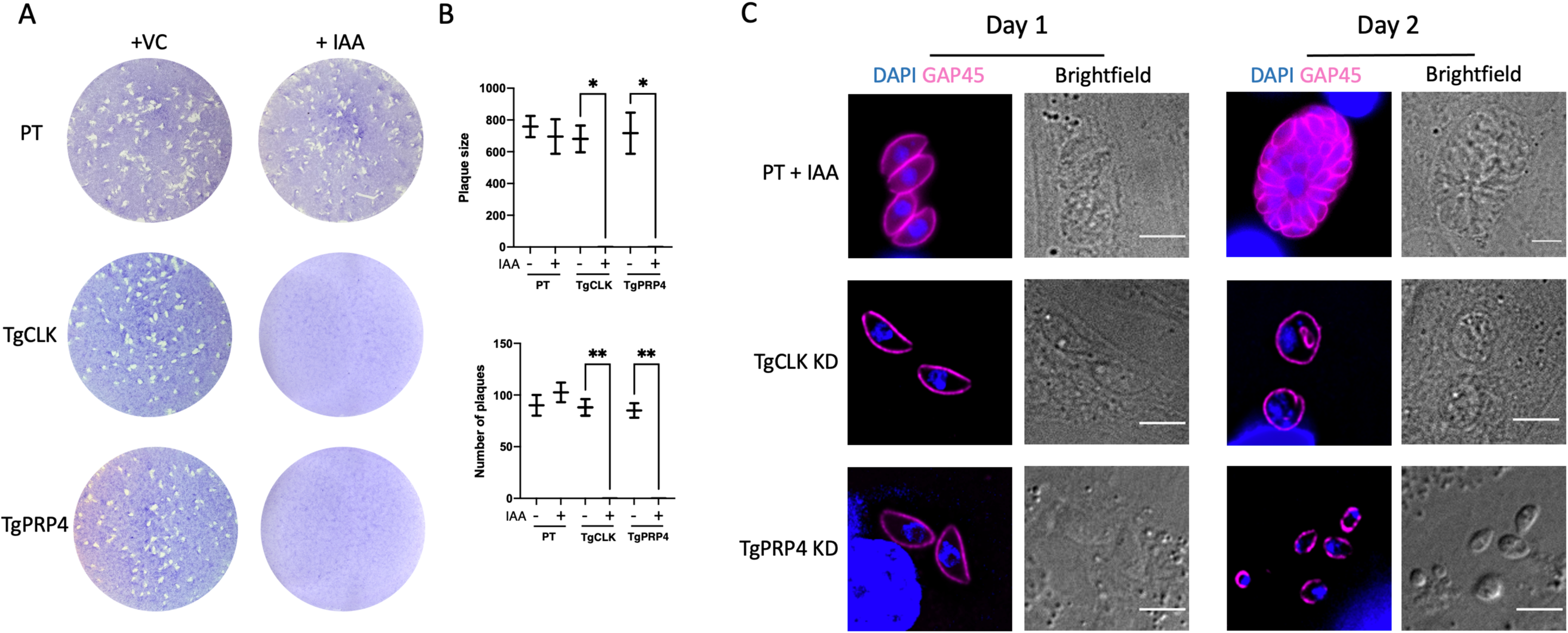
**Growth of ablated AID-HA tagged parasites.** (A) Representative plaque assay images of AID-HA tagged parasites in HFF cell monolayers stained with crystal violet. (B) Quantification and statistical analysis of the mean size (top) and number (bottom) of plaques from (A) in 2 biological replicates using the Student’s paired t-test. *, P< 0.05; **, P< 0.01. (C) Representative time-course immunofluorescence microscopy images of intracellular AID-HA tagged parasites following the addition of IAA. The parental line was used as the negative control in each (A-C) assay.

### Ablation of *Tg*SRs, *Tg*CLK and *Tg*PRP4 perturbs gene expression

To determine the cause of the defects, and whether the putative *Tg*SR proteins and kinases are alternative-splicing factors, we performed poly(A) selected RNA-seq on the parental and mutant lines. We sequenced cDNA from the parental lines, the *Tg*SR direct knockout mutants, and the *Tg* kinase knockdown mutants after incubation with 500 µM IAA or EtOH for 8 hours, in biological triplicates. The choice to assay the parasite RNA at the 8-hour time point was based on our previous work on *Tg*SR3, which revealed the greatest change in AS 6-8 hours post-induction with minimal pleotropic effects (30). We also sequenced the RNA from the parental line with and without IAA as the control group, to test the indirect effect of IAA on the parasite transcriptome. We sequenced the 30 samples on the Illumina NovaSeq6000 platform and obtained an average of ~39 million paired reads (2 x 150 bp) per sample. Reads were analysed for quality with FASTQC and mapped to the *T. gondii* genome.

We first investigated whether the transcriptional profiles of each sample were similar or different from each other using multi-dimensional scaling (MDS) plots as created in Limma (42). MDS plots allow us to observe transcriptional changes in an unsupervised manner and identify sample clusters, be it from biological groups or batch effects. The MDS plot of the *Tg*SRKOs samples revealed clustering by their respective biological groups distinct from the parental line (Fig 5A top). Similarly, the *Tg*CLK and *Tg*PRP4 knockdown samples clustered within their respective biological groups separate from the vehicle controls (Fig 5A bottom). The vehicle controls clustered together with all the parental lines-uninduced, with IAA or with EtOH. These indicate that simply adding IAA or tagging the genes does not alter the transcriptome profiles of the parasites, while depletion of the *Tg*SR proteins or kinases do result in changes in gene expression. Moreover, the changes appear to be distinct between parasite lines, and consistent between replicates.

**Figure 5.**
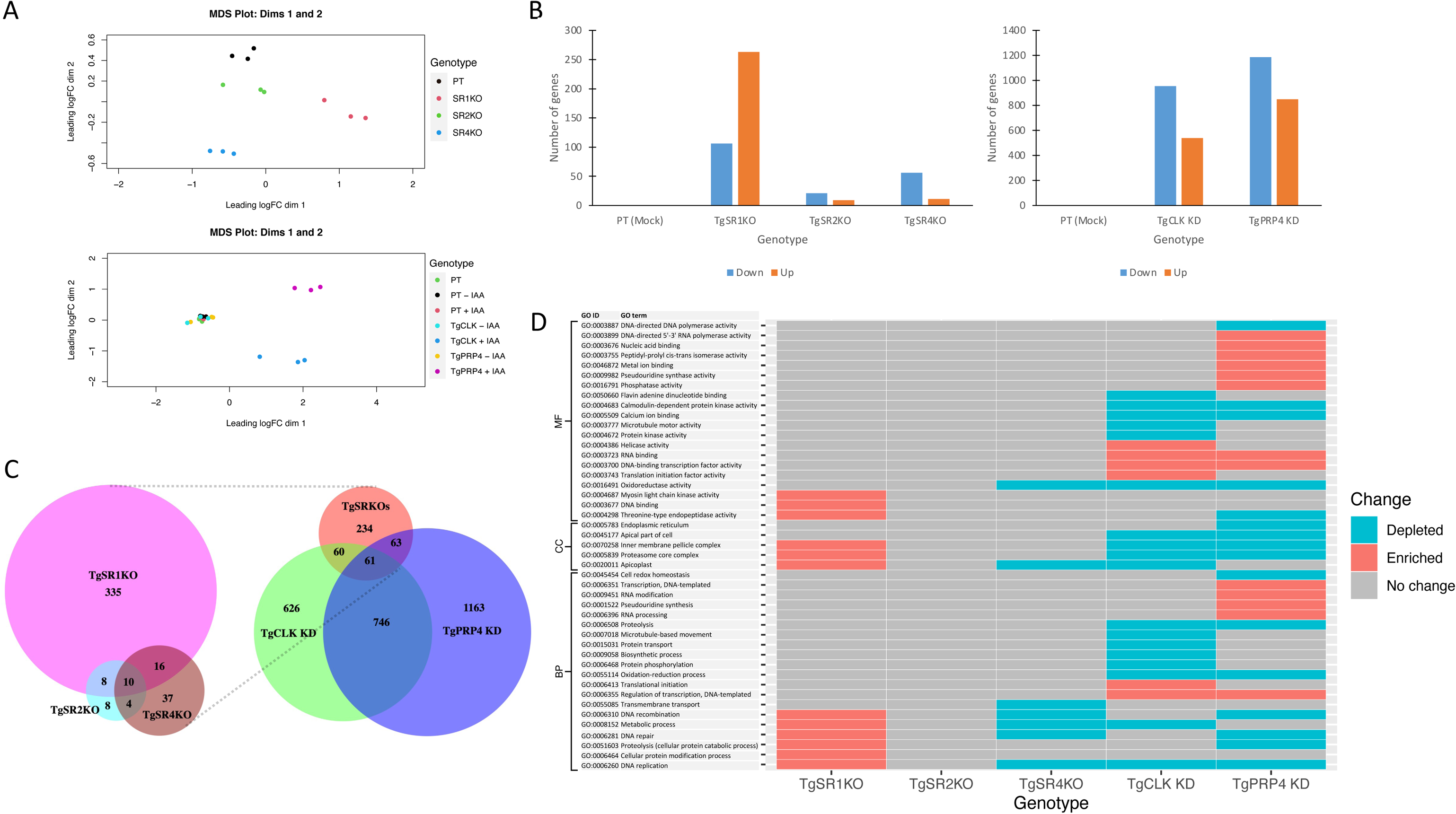
**Differential expression analysis of genes for ablated *Tg*SR and putative *Tg*SR kinase parasites.** (A) MDS plots of log-CPM values for each parasite sample. Distances represent the leading fold-change for the 500 genes most divergent between each pair of samples. (B) Bar graphs of differentially expressed genes that were downregulated (down) or upregulated (up). Only genes with an adjusted p-value of ≤ 0.05 and fold change of ≥ 2 were considered differentially expressed. (C) Proportional Venn diagram showing the overlap of differentially expressed genes between each parasite genotype/condition. (D) Heatmap of GO terms that were over-represented in upregulated (enriched) or downregulated (depleted) genes for each parasite genotype/condition as identified using GSEA. Only terms with a p-value of ≤ 0.05 and FDR of ≤ 0.25 were considered over-represented.

We then determined the exact genes that were statistically significantly differentially expressed using the linear modelling approach of Limma (42). We set a significance threshold of 0.05 or less for the adjusted p-value, and required a minimum log-fold change of at least 2 for all comparisons. As expected, there were no genes that were significantly changed in the parental line with or without IAA control group, but many in the mutant lines (Fig. 5b). There were hundreds of differentially expressed genes in *Tg*SR1KO (Up-263; Down-106), and tens for *Tg*SR2KO (Up-9; Down-21) and *Tg*SR4KO (Up-11; Down-56) parasites. More remarkably, thousands of genes were differentially expressed for *Tg*CLK-KD (Up-540; Down-953) and *Tg*PRP4-KD (Up-848; Down-1183). Generally, there were more genes that were downregulated than upregulated, with the exception of *Tg*SR1KO, which exhibited the opposite. This is particularly obvious on the mean-different (MD) plots as presented in Figure S2, which further showed that the magnitude of differential expression for transcripts was not strongly influenced by average transcript abundance. Consistent with the MDS analysis, the differentially expressed genes for each parasite line did not strongly overlap. One-to-one comparisons showed only around 50% or less overlap (Fig. 5C). For the *Tg*SRKOs, only 10 DE genes were common between the 3 lines. Between the *Tg*SRKOs, *Tg*CLK, and *Tg*PRP4, only 61 DE genes were common. It should be noted that each of the three *Tg*SR transcripts were identified as significantly downregulated in their respective KO lines. Additionally, IGV snapshots showed no reads mapping to the excised gene locus for *Tg*SR1KO and *Tg*SR4KO, and altered mapping for *Tg*SR2KO (Fig. S3). These further confirm that the genes were successfully knocked out.

To understand what processes the differentially expressed genes might be involved in, we performed gene ontology (GO) enrichment analyses using GSEA (43). GSEA is able to identify GO terms that are over-represented in upregulated (enriched) or downregulated (depleted) genes. We summarised the results in a combined heatmap (Fig. 5D) that included all significantly (p ≤ 0.05, FDR ≤ 0.25) over-represented terms for each mutant/condition. The analysis revealed similar GO terms that were over-represented in *Tg*SR1KO and *Tg*SR4KO. In particular the terms “DNA recombination”, “DNA repair”, “DNA replication”, and “metabolic process” were enriched in differentially expressed genes. These pathways have been previously linked with SR proteins (44) (see discussion). The GO term with the highest normalised enrichment/depletion score (NES/NDS) for both mutants was the cellular component “apicoplast”, though the reason for this is unclear. Intriguingly, the terms were enriched in *Tg*SR1KO, but depleted in *Tg*SR4KO, suggesting some complementary roles between the two *Tg*SR proteins. Another enriched category in the *Tg*SR1KO parasites was proteolysis. Potentially, abnormal splicing caused by the *Tg*SR1 ablation was producing increased levels of aberrant proteins that required higher levels of degradation. Unsurprisingly, no GO terms were over-represented in *Tg*SR2KO due to the low number of DE genes.

Compared to the *Tg*SRKOs, more GO terms were significantly over-represented for the two kinase knockdowns (*Tg*CLK-24; *Tg*PRP4-29). The GO assignments were quite varied, with almost equal weighted distribution between the three categories of “molecular function (MF)”, “cellular component (CC)”, and “biological process (BP)”. While there were some overlaps of GO terms, the general profiles were distinctive, suggesting some divergent roles between the two genes. Terms with high NES/NDS and overlapped included terms for or related to RNA binding/processing, DNA transcription, and proteolysis. These terms are concordant with a disrupted splicing process that was impacting immediate downstream/upstream processes. We noted that *Tg*PRP4-KD distinctly affected the “RNA modification” pathway, particularly in relation to “pseudouridine synthesis”/ “pseudouridine synthase activity”. Similar to *Tg*SR1KO and *Tg*SR4KO, *Tg*PRP4-KD also affected “DNA recombination” and “DNA repair”. On the other hand, *Tg*CLK-KD had more protein related terms, such as “protein phosphorylation”, “protein transport” and “translation”. Together, these results indicate that the genes are involved in the splicing process, but that they are also more widely implicated in other cellular functions as has been previously reported of kinases of SR protein (45).

### Ablation of *Tg*SRs, *Tg*CLK and *Tg*PRP4 perturbs alternative splicing

Because the genes we disrupted are predicted to be alternative splicing (AS) factors, we sought to determine the perturbation to AS in these parasites. We use the term AS to describe a splicing deviation from the annotated canonical model, but the distinction between constitutive and alternative splicing is not clear even in the literature (15). Regardless, there are several strategies that are commonly applied for differential splicing (DS) analyses. The more widely-used ones include a subgenic feature count-based method (e.g., DEXSeq (46), edgeR (47) and JunctionSeq (48)), or an event/junction-based method (e.g., dSpliceType (49), MAJIQ (50), and rMATS (51)). While the latter is able to more easily identify and classify types of alternative splicing changes, it suffers from less robust statistical power that is afforded by the former methodology (52). Relatively recently, a program called ASpli (53) was developed. ASpli combines both approaches within a single tool by utilising the methodology of DEXSeq, the statistical framework of edgeR, and additional evidence from junction inclusion indexes to robustly identify the types and changes in AS. We used ASpli to analyse our samples and identified significant changes in AS events corresponding to many genes for our KO/KD parasites, but none in the control group. Of the *Tg*SR knockouts, *Tg*SR1KO resulted in the greatest number of changes (472 events; 328 genes), followed by *Tg*SR4KO (110 events; 68 genes) and *Tg*SR2KO (80 events, 52 genes). Though notable, these mutants produced far fewer changes than knocking down *Tg*CLK or *Tg*PRP4, which resulted in widespread AS changes (*Tg*CLK-7081 events; *Tg*PRP4-13137 events) that affected thousands of genes (*Tg*CLK-2786 genes; *Tg*PRP4-3713 genes). The numbers represent 53.33% and 71.76% of multi-exon transcripts that were reliably detected in the *Tg*CLK and *Tg*PRP4 samples respectively, indicating their extensive role in regulating splicing.

We then assigned AS events to the 4 main AS types: intron retention (IR), alternative 5’ splice site selection (Alt5’SS), alternative 3’ splice site selection (Alt3’SS) or exon skipping (ES). We looked at the types of differential AS events in our data sets and found that IR events occurred at the highest frequency across all comparisons (Fig. 6A top). This was followed by Alt5’/3’SS and ES in the *Tg*SRKOs, consistent with previous findings of other SR proteins (26, 54). In contrast, ES was the second most common change when *Tg*CLK or *Tg*PRP4 was depleted, followed by Alt5’/3’SS. The AS events corresponded proportionally to gene numbers (Fig. 6A bottom). We looked at the productivity of the over-represented intron retention events, as defined by whether the event introduced a premature stop codon, and find that virtually all (>99%) events would lead to premature termination. This implies that most of the AS events were unlikely to translate to new protein isoforms. Prior studies have shown that SR proteins function by directly enhancing and/or repressing splicing (24). To determine if the *Tg*SR proteins or kinases preferentially activate or repress splicing, we visualised the changes of intronic events as volcano plots (Fig. 6B). *Tg*SR1KO increased and decreased the AS events at almost equal proportions, indicating a dual role. Conversely, knocking out *Tg*SR2 and *Tg*SR4 predominantly decreased levels of intron retention, indicating their primary role as splicing repressors.

**Figure 6.**
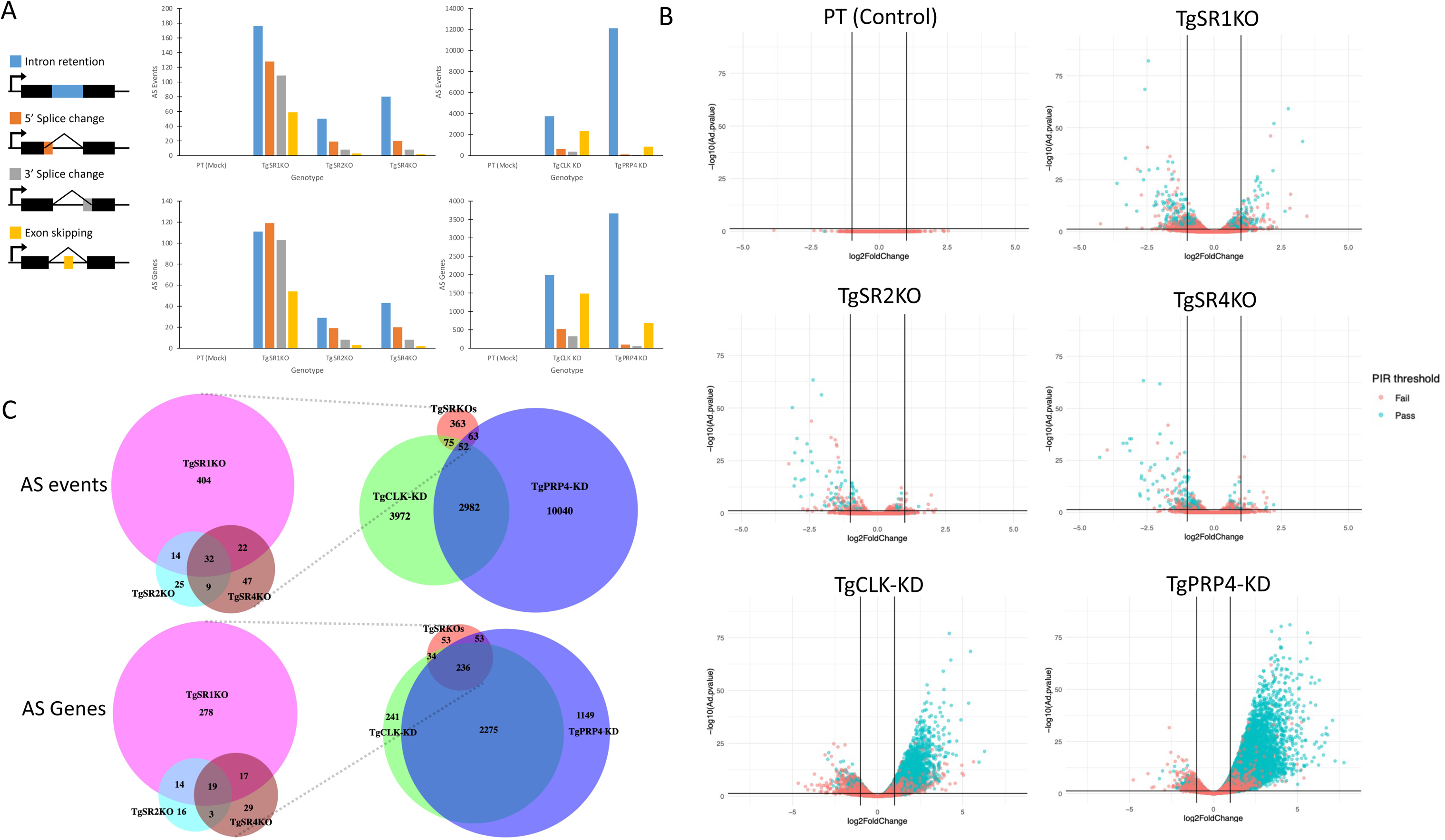
**Differential splicing analysis of junction and genes for ablated *Tg*SR and putative *Tg*SR kinase parasites.** (A) Bar graphs of differentially spliced junctions (top) and genes (bottom) for each major type of alternative splicing. Only AS junctions/genes with an adjusted p-value of ≤ 0.05, fold change of ≥ 2 and difference exceeding 10% were considered differentially spliced. (B) Volcano plots of differentially spliced introns. Percent intron retention (PIR) is a junction inclusion index that represent the proportion of intron retention events to the canonical junction. The PIR threshold represents whether the proportion equal or exceed 10% in difference and 2-fold change, which we use as an additional level of evidence to support changes in AS. (C) Proportional Venn diagram showing the overlap of differential AS junctions (top) and genes (bottom) between each parasite genotype/condition.

Strikingly, most of the AS events that were detected for *Tg*CLK and *Tg*PRP4 KD increased dramatically. This observation implies that the two kinases are strong repressors of AS, or that they are simply required for normal splicing, though it is not clear if this is mainly through the SR proteins. To understand the phenomenon better, we looked at the overlap of differential AS events between the parasites, reasoning that we would expect a stronger connection for the proteins that overlapped in function or were co-dependant. In line with the partial redundancies identified of previously-studied SR proteins in other organisms (15), there is moderate (~50%) one-to-one overlap between the three *Tg*SR proteins for differential alternative splicing (whether measured at a per-event or per-gene basis) (Fig. 6C). Interestingly, there is a strong (85.90%) overlap between the *Tg*SRs and the kinases that we originally hypothesized might regulate those SRs, but only when considered at a gene level, and not at a per-junction level. Despite *Tg*CLK-KD and *Tg*PRP4-KD having thousands more differential AS events, only 34.36 % (190/553) of *Tg*SRKOs differential AS events overlapped with any of the two kinases. These data argue against a simple relationship where one SR is regulated by one SR-regulating kinase. The high level of gene overlap can be partially attributed to the large proportions of multi-exon genes that had differential AS events in *Tg*CLK-KD and *Tg*PRP4-KD, but also points to these kinases regulating splicing through mechanisms that extend beyond the SR genes we studied.

To explore other mechanisms regulated by *Tg*CLK-KD and *Tg*PRP4-KD, we looked at whether particular GO terms were enriched for our differential AS genes, as we did for the DE genes. The results are presented in supplementary Figure S4. Of the *Tg*SRKOs, only *Tg*SR1KO had GO terms which were significantly enriched. The terms were all only categorised as BPs and were rather broad without clear connections. For *Tg*CLK-KD and *Tg*PRP4-KD, multiple GO terms were assigned as well though for *Tg*CLK-KD, they were mostly related to “DNA replication”. Of note, the most significantly enriched term-“helicase activity”, was also highly enriched in the DE analysis. Such connections are less clear for *Tg*PRP4-KD.

### Does alternative splicing impact transcript abundance?

Previous studies have established the ability of SR proteins to regulate abundance of individual transcripts through direct AS targets, or through AS independent mechanisms like transcription elongation and mRNA export (19). To determine how the differential AS events might be impacting gene expression, we first looked at whether the differential AS genes corresponded with the genes that we had identified as being differentially expressed. We find that the differential AS genes and differentially expressed genes overlapped poorly, with the exception of *Tg*PRP4-KD (Fig. S5). For the *Tg*SRKOs, the overlap is only around 10% or less. *Tg*CLK-KD has slightly better overlap (~20%) and *Tg*PRP4-KD the highest (~30%), though the results could again be attributed to the high proportion of genes that were implicated in each subset. We looked at the expression of genes with differential AS events using hierarchical clustering as plotted on a heatmap, and found that while the protein/gene depleted samples did cluster separated from the control samples, the exact differences were subtle in most cases (Fig. S5). Thus, our findings indicate that the SR proteins and kinases were predominantly impacting gene expression through downstream AS targets (e.g. transcription factors or compensatory pathways) or AS independent mechanisms. Regardless, the data is not consistent with AS having a major role in driving regulation of transcript abundance. It can be noted the poor overlap is also indicative of the ability of ASpli in detecting differential AS events that is not confounded by changes in transcript level.

### What are the SR proteins binding to?

In general, SR proteins typically bind degenerate purine-rich sequences on exons known as exonic splicing enhancers (ESE). We set out to determine if such elements were present in the differential AS events by using multiple expectation-maximization for motif elicitation (MEME) (55) on the region 150 bases upstream and downstream of the 5’ or 3’ splice site. We focused on IR events given their overrepresentation, and used matched number of junctions with no change in AS as a comparison. The results are summarised in Figure 7. The search yielded the motif of GRAGRAA (R = G or A) for the 5’ splice site, and GMGGARVAR (M= A or C; V = A, C or G) motif for the 3’ splice site, that were significantly enriched in *Tg*SR1KO. These purine rich motifs closely match the previously confirmed SRSF1-binding motif core GAAGAA, characterised in human cells (56) which is known to be degenerate and context sensitive (57). To understand where these motifs might be localised in the sequence, we graphed the positional probability frequency plots of the motifs. Consistent with the characteristics of ESEs, the graphs showed a spike of the motifs near the exonic region of the splice junction, followed by a decline within the intronic region. Similar purine rich sequences were enriched in *Tg*SR3KO. However, these were much less defined, and the spike of the 5’ site motif occurred within the intron rather than exon. The position of SR protein binding within introns is further indicative of a splicing repressor role (24), consistent with the results from our splicing analysis. No motifs were significantly enriched for *Tg*SR2KO.

**Figure 7.**
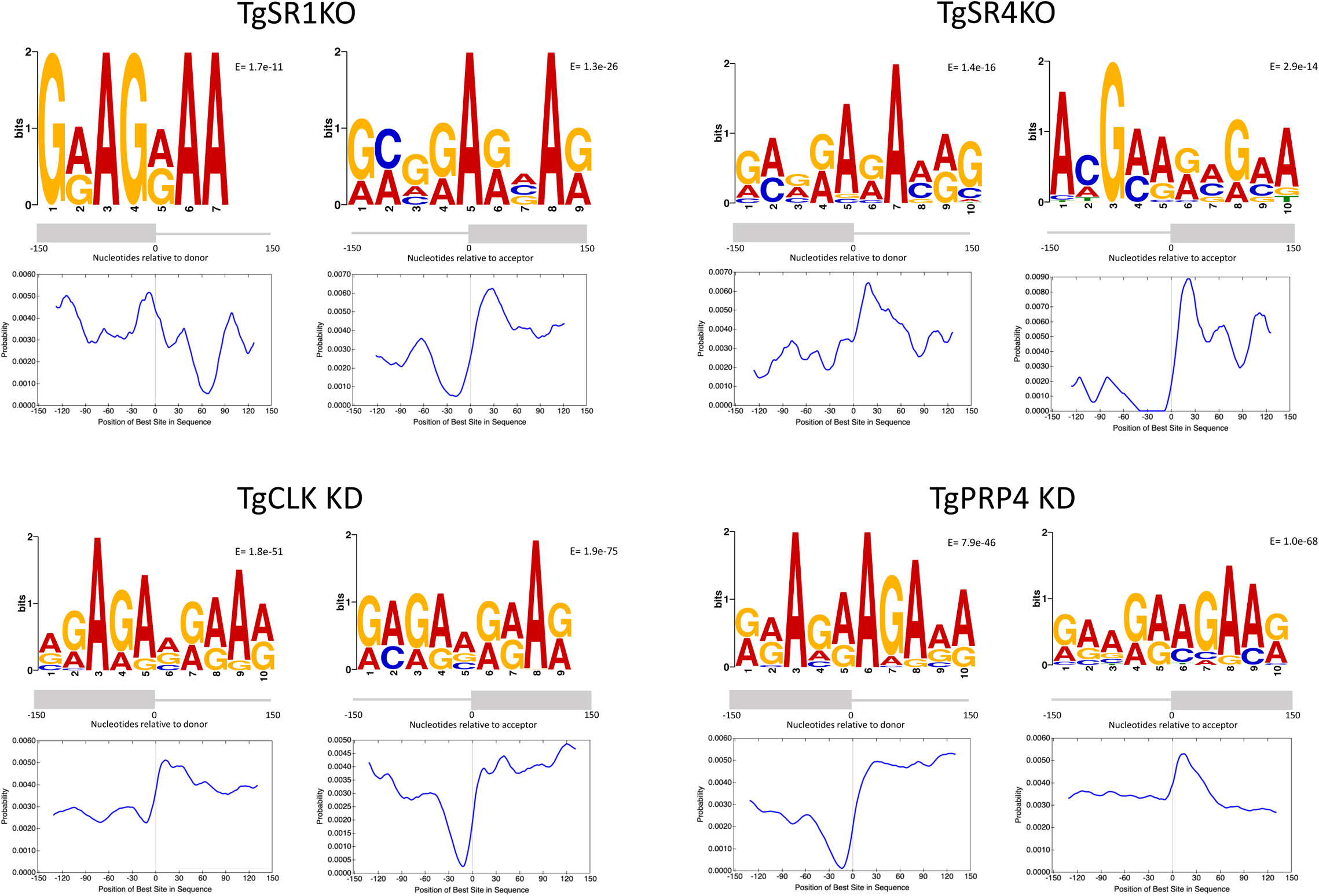
**Sequence logos for the motifs enriched** 150 bases upstream of and downstream from the splicing acceptor (3′ splice site) and splicing donor (5′ splice site) of differential intron retention junctions for each parasite genotype/condition, where present. The positional probability frequency plots of the motifs are shown below each of the motifs.

Although *Tg*CLK and *Tg*PRP4 were not predicted to directly bind RNA, we further investigated their function in modulating AS by repeating the MEME analysis. The results reveal multiple purine-rich sequences that were enriched in each case. We present the most significant ones in figure 7. In the case of *Tg*PRP4, the motif GAAGAAGAA could be discerned from the most significantly enriched 3’ site sequence. The sequence is a well-known ESE motif in humans and plants (58, 59), and matches the SRSF1 binding motif core. Other motifs are less specific. In addition, probability frequency plots of the motifs were quite varied. Generally, a spike was observed around the splice sites, but this occurred within both introns or exons. In some cases, rather than a spike, the plot reveals a depletion of the motif around the splice site instead. Taken together, these results suggest that *Tg*CLK and *Tg*PRP4 are indeed modulators of widespread AS events, though the mechanism is unclear.

## Discussion

RNA splicing is integral in the transcriptional and post-transcriptional control of genes in metazoans but less is known in apicomplexan parasites. Recent transcriptomic studies have revealed that like metazoans, widespread alternative splicing (AS) occurs in many unicellular parasites, contributing to varied processes such gene expression, protein trafficking, and antigenic diversity (60). SR proteins and kinases that phosphorylate SR proteins are modulators of AS that have been partially characterised and validated as drug targets in *Plasmodium* (27–29). However, less is known about these genes in *T. gondii* (30). In this study, we showed that the previously uncharacterised putative *Tg*SR proteins are abundantly expressed in the parasites and are found localised to nuclear speckles. Nuclear speckles are sites of splicing factors storage and assembly, and all SR proteins have been found localised to this structure (21). In metazoans, nuclear speckles are normally stable during interphase and disassemble/become disperse during cell division (61). Subsequently, nuclear speckle proteins form mitotic interchromatin granules (MIGs) within the cytoplasm before relocalising to nucleus at the end of mitosis (61, 62). Interestingly, our results show that nuclear speckles are stable even during mitosis in *T. gondii*. This is not surprising given that mitosis in apicomplexan parasites occurs without breakdown of the nuclear envelope and little condensation of chromatin (63). This does however highlight an appreciable difference between metazoan and *T. gondii* SR proteins and perhaps other splicing factors that has not been explored.

Congruous with the localisation data, all three characterised *Tg*SR genes impact mRNA splicing. When knocked out, AS is disrupted, though to different extents in the three *Tg*SR proteins. This has been similarly observed for the suite of SR proteins identified in other organisms (54), and for the overexpression of *Tg*SR3 in *T. gondii* (30).Previous analyses have revealed that the activity of SR proteins in modulating AS correlates positively with the number of arginine-serine (RS) repeats that the SR protein possesses (64). Consistent with this, *Tg*SR1 has the highest molecular weight and number of RS repeats, and governs the greatest number of AS events. Comparatively, *Tg*SR2 and *Tg*SR4 are much smaller and exerted a more modest effect on AS. We also observed some overlap of affected AS events and genes between the three *Tg*SR proteins. This further mirrors the partial redundancies identified of other SR proteins. For example, several human SR proteins are able to exert the same splicing pattern on pre-mRNA (17). It is thus not surprising that individual deletions of the *Tg*SR proteins do not seriously impact parasite survival. In *C. elegans*, only simultaneous RNAi of all, and not individual SR proteins, results in embryonic lethality (65). The RNAi pathway is deficient in apicomplexan parasites (66) and so more complex combinatorial deletions of *Tg*SR proteins would be of interest in future experiments.

Notwithstanding the non-essentiality of individual SRs, different SR proteins have distinct functions and their specific activity is well established. Typically, the RRM domain of SR proteins binds to purine rich exonic sequences on pre-mRNA known as exonic-splicing enhancers (ESEs) and promote splice-site selection through direct interactions with other RS domain containing splicing factors that form the early spliceosomal complex (15, 67, 68). However, SR proteins may also bind to intronic regions and suppress splicing (54, 69–71). This is less commonly seen, and it is not exactly clear how these SR proteins inhibit or alter recruitment of the spliceosome complex. The deletion of *Tg*SR2 and *Tg*SR4 almost exclusively decreased subsets of intron retention events that were not linked with any specific pathways, implying a general splicing repressor role, like the human SRSF10 protein (69). This is further supported by the intronic purine rich sequence that was enriched in *Tg*SR4 differential intron retention events. On the other hand, *Tg*SR1 appears as a more conventional SR protein. In our previous phylogenetic analysis, *Tg*SR1 was most closely related to SRSF1 (30). Similar to SRSF1 (72), *Tg*SR1 enhances and represses AS events, and exonic purine-rich motifs were enriched in the AS junctions. The motifs matched the ESE core motif for SRSF1, which supports some evolutionally conservation between the two genes. We note, however, that SR protein RNA interactions defy easy categorisation due to many factors including motif degeneracy, competition with other RNA binding proteins and dynamic RNA secondary structures (73). Even in the well-studied mammalian splicing machinery system, SR protein-RNA binding specificities remains to be fully characterised (24).

Intriguingly, while the *Tg*SR genes deletions caused changes in gene expression, the changes were not directly linked to AS events. SR proteins are known to regulate gene expression through multiple pathways. In one pathway, SR proteins have been shown to directly interact and influence components of gene transcription. For example, SRSF2 is able directly regulate the elongation rates of RNA polymerase II (74). In other cases, specific SR proteins appear to play a role in mRNA export from the nucleus and translation. SRSF3 and SRSF7 have been shown promote the export of intronless RNAs (75), and SRSF10 repress translation via interaction with the peptidyltransferase centre of 28S rRNA, a mechanism thought to inhibit the differentiation of primary neuronal cells (76). An important but less direct pathway of gene expression regulation is through the nonsense-mediated decay (NMD) pathway. SR mediated AS often generate RNA isoforms that are inherently unstable/non-productive due to pre-mature terminating codons (PTCs) encoded in retained introns or translational frameshift events that target the mRNA for degradation through the NMD pathway (77). This particular pathway has been implicated in the regulation of stem cell differentiation (78) and may be similarly involved in the sexual differentiation of *P. berghei* (79). Our data showed that the SR linked AS events predominantly resulted in intronic PTC containing transcripts, which would presumably create targets for the NMD pathway. However, the lack of gene expression changes linked with the AS events does not support this indication. While the discrepancy has also been observed for *Drosophila* SR proteins (54), it is nonetheless unusual because the AS transcripts will likely affect gene translation, be it the production of altered or truncated deleterious proteins (80). The over-represented GO terms relating to proteolysis in *Tg*SR1 is consistent with this hypothesis. We do not see this for *Tg*SR2 or *Tg*SR4 because the ablations decreased, rather than increased non-productive splicing.

In addition to splicing or gene expression, SR proteins have been implicated in other roles. For example, depletion of SR protein SF2/ASF in chicken DT40 cells, and SC35 in mouse embryo fibroblasts induces DNA damage and recombination (81, 82). This link with genomic instability is likely due to a defect in transcriptional elongation that results in exposed ssDNA susceptible to nucleases or modification (44). Our GO analysis reveals that DNA repair and recombination were similarly engaged when *Tg*SR1 and *Tg*SR4 was ablated. This further reinforces the multi-faceted nature of SR proteins.

Perhaps most interestingly, our results identified *Tg*CLK and *Tg*PRP4 as extensive regulators of mRNA splicing, but the splicing events overlapped poorly with that regulated by the three *Tg*SR proteins. In humans, the localisation and functions of SR proteins are extensively regulated through phosphorylation on serine residues of the RS domain mainly by members of two kinase families-SRPK and CLK, in a “relay” type process (83). The process is crucial for the assembly of spliceosomal components and for AS to occur (84). In *T. gondii,* previous global phosphoproteomic data have revealed that the N-terminal RS domain of TgSR proteins are extensively phosphorylated on serine residues (85). However, the presence of relatively fewer kinases of SR proteins suggests some evolutionary divergence from metazoans. The poor overlap of AS events regulated by the *Tg*SR proteins and the two kinases further highlights this. One caveat is that we did not investigate *Tg*SR3, which we had previously characterised through overexpression experiments (30). We attempted to ablate *Tg*SR3 here as well but this was unsuccessful. Further, we cannot conclusively identify the exact influence of the kinases on the *Tg*SRs due to the inherent redundancies and competition of SR proteins. It is possible that ablating a combination of the *Tg*SR proteins will result in a greater overlap. However, even the sum of combined AS disruptions from the *Tg*SR protein deletions is dwarfed by the extensive splicing changes from depleting either of the kinases. Moreover, the depletions affected the majority of multi-exon genes, rather than a distinct subset. Given that the *Tg*SR proteins also differed in their specific function, the data suggests that the kinases were predominantly influencing splicing through other factors.

Members of the kinase families may be predominantly known to phosphorylate SR proteins, but many of the kinases also have distinct roles outside of phosphorylating SR proteins. For example, human SRPK2 is involved in phosphorylating Prp28, an RNA helicase, in the spliceosome assembly process. Our GO analyses similarly revealed that *Tg*CLK depletion impacted helicase activity, which would in turn affect splicing and gene expression. We also showed that ablation of *Tg*PRP4 impacted the pseudouridine RNA modification pathway, particularly pseudouridine synthesis, which is essential for gene expression regulation and pre-mRNA splicing, as is observed in Xenopus oocytes (86), and humans (87). A further complication is that kinases that phosphorylate SR proteins directly interact one another and other splicing factors. Human SRPK1 for example has been shown to control the release of CLK1 from SR proteins (88), and PRP4 directly interacts with U5 snRNP, a component of the spliceosome (89). Nonetheless, our results indicate that while individual SR proteins would appear to be poor drug targets, inhibitors to *Tg*CLK and *Tg*PRP4 would simultaneously disrupt many genes and pathways, killing the parasites. This is supported by the recent characterisation of a *Plasmodium* splicing-regulating kinase CLK3 (the homologue of *Tg*PRP4 in this study), which was validated as a target for potent and specific drugs (29).

SR proteins and kinases that phosphorylate SR proteins are multifaceted and defy easy characterisation of their function. Our study aimed to elucidate the AS framework of *T. gondii,* through the broad characterisation of all these proteins. Using a transcriptomic approach, we revealed that *Tg*SR proteins do mediate subsets of AS splicing and caused changes in gene expression. Similarly, two putative kinases of the SR proteins, *Tg*CLK and *Tg*PRP4, are crucial mediators of the splicing process and are essential to parasite survival. However, SR proteins are only required to mediate a relatively small subset each of AS events and genes, while *Tg*CLK and *Tg*PRP4 are essential for the splicing process. These findings shed some light on the molecular actors that mediate transcriptional and post-transcriptional regulation and also highlight the complex nature of transcript regulation which requires further dissection.

## Methods

### Parasite cultures and manipulation

Prugniaud Δ*hxgprt* Δ*ku80* (PruΔku80) and subsequently derived parasite lines were cultured on confluent monolayer human foreskin fibroblasts (HFFs) in Dulbecco’s Modified Eagle medium (DME) supplemented with 1% v/v fetal calf serum (FCS) and 1% v/v Glutamax at 37 °C in a humidified 10% CO_2_ incubator. Transfections were performed on the Nucleofector 4D system (Lonza) using 2 × 10^6^ tachyzoites in 20 μL of buffer P3 (Lonza) with the F1-115 pulse code, followed by selection in either 1 μM pyrimethamine or mycophenolic acid and xanthine. Clonal parasite lines were obtained by limiting dilutions onto HFFs in 96-well plates and selecting parasites from single plaque wells for propagation. We performed plaque assays by inoculating 200 tachyzoites onto 6 well plates of HFFs and allowing the parasites to grow undisturbed for 8 days. The plates were then fixed in 80% v/v ethanol and stained with crystal violet (Sigma). Images were scanned and analysed with FIJI (90) using the ViralPlaque macro (91). Averages over replicates were tabulated and statistically analysed with a paired t-test. Bradyzoite differentiation was done by inoculating tachyzoites onto HFFs at a multiplicity of infection (MOI) of 0.1 in RPMI-HEPES (pH 8.1-8.2) supplemented with 5% v/v (FCS) at 37 °C in a humidified incubator (without CO2). The media was refreshed every 2 days to maintain pH for 3 times prior to further assays.

### DNA and Plasmid construction

Genes of interests were edited using a previously established CRISPR/Cas9 approach for *T. gondii* (32, 33). Briefly, a sgRNA expressing plasmid was co-transfected with a targeted insert sequence for each modification. We used EuPaGDT (92) to design the CRISPR guide and Q5 mutagenesis (NEB) to clone the sequence into the pU6-Universal (a gift from Sebastian Lourido-AddGene #52694) plasmid. Insert sequences were obtained via PCR of the following plasmids: pLIC-3xHA-HXGPRT (93) for HA tagging, pLoxP-DHFR-mCherry (Addgene #70147) (36) for gene knockout, and mAID-3XHA-HXGRPT-TIR1 for AID tagging. We used Gibson assembly of the pTUB1: OsTIR1-3FLAG (Addgene #87258) (94) and pTUB1: YFP-mAID-3HA (Addgene #87259) (94) plasmids to create mAID-3XHA-HXGRPT-TIR1. 40 bp homology flanks of the insertion site was directly incorporated to the insert sequence via the PCR primers. Primers are listed in Supplementary File S1.

### Phylogenetic analyses

Sequences of the human 3 SRPKs (SRPK1–3), 4 CLKs (CLK1–4), PRP4 and DRYK1a kinases were extracted from Genbank (95) and used to identify homologues in *T. gondii, P. falciparum, pombe* and *A. thaliana* using BLASTp (95) in ToxoDB (96), PlasmoDB (38) and the default non-redundant NCBI databases (97). Alignment, curation, and phylogenetic tree building was done using the Phylogeny.fr (98) web service with default options.

### IFA and Western blotting

IFAs-*T. gondii*-infected host cells on coverslips were fixed in 4% v/v paraformaldehyde in PBS for 10 mins, permeabilized in 0.1% v/v Triton X-100 in PBS for 10 minutes, and blocked in 3% w/v BSA (Sigma) in PBS for 1 hr. We also performed primary and secondary antibody incubations at room temperature for 1 hr, with three PBS washes after each step. For primary antibodies, we used the rabbit-αGAP45 (99), rat-αHA 3F10 (Roche) and rabbit-αSRS9 (100) antibodies at a 1:1000 dilution, and Rhodamine-Dolichos Biflorus Agglutinin (Vector Labs) at a 1:200 dilution, in blocking buffer. For the secondary antibodies, we used Alexa Fluor 488, 594 or 647 (Thermo Fisher) at a 1:1000 dilution in blocking buffer. 5 μg/ml DAPI in PBS was used to stain the nucleus in the penultimate wash before mounting onto microscope slides with Vectashield (Vector Labs). Images were taken with either the DeltaVision Elite or the DeltaVision OMX Blaze microscopes.

Western blotting-Freshly egressed (90%) HA-tagged or direct knockout parasites lines from single T25 flasks were harvested and lysed in 50 μl lysis buffer (1% v/v Triton-X 100, 1 mM MgCl2, 1X cOmplete EDTA-free protease inhibitor cocktail (Sigma), 0.1 % v/v Benzonase (Merck)) on ice for 30 mins. AID-tagged parasite lines were lysed similarly, but we only used intracellular cellular parasites released by scrapping and passing infected HFFs through 27-gauge needles. Samples were then mixed with equal volume of loading buffer (2 × Laemmli buffer, 5% (v/v) β-mercaptoethanol) and boiled for 10 mins. We ran the samples on a 4%–15% SDS-PAGE gel (BioRad), and transferred the proteins onto a nitrocellulose membrane. Blocking and antibody incubations were done for 1 hour in 5% milk in PBS-T (0.05 % Tween-20 in PBS). We used the αHA and αGAP45 primary antibodies as used for IFAs, and horseradish peroxidase-conjugated goat anti-rat antibody (Santa Cruz Biotechnology) diluted 1:2000 as the secondary antibody. Membranes were washed with PBS-T 3 times after each step and finally imaged on the ChemiDoc MP Imaging System (Biorad).

### Library prep and RNA-sequencing

Prior to RNA extraction, parasites were allowed to infect HFF monolayers for 48 hours before the cells were harvested. AID-tagged parasites were treated with EtOH or IAA for 8 hours before collection. The choice to assay the parasite RNA at the 8-hour time point was based on our previous work on *Tg*SR3, of which overexpression caused the greatest changes in alternative splicing 6-8 hours post-induction, and subsequent pleotropic effects manifesting at later time points. The parasites were mechanically released from HFF cells by passing the samples through a 27-gauge needle 3 times and pelleted. Parasite RNA was extracted using TRI Reagent (Sigma) followed by the RNeasy MinElute Cleanup Kit (Qiagen) according to the manufacturers’ protocol. DNase 1 (Qiagen) was used to remove contaminating DNA followed by a second cleanup. We used the services of Victorian Clinical Genetics Services (Melbourne) for further QC (Agilent TapeStation), poly-A enrichment, paired-end 150 bp cDNA library construction using the TruSeq stranded mRNA kit (Illumina) and sequencing on the NovaSeq 6000. All RNA-seq data have been deposited in the NCBI Sequence Read Archive BioProject under accession number PRJNA738301.

### Transcriptome analyses

Sequencing data was first checked for quality with FastQC (v.0.11.7). We then utilised STAR (v.2.7.5b) (101) to align sequencing reads to the parasite genome from ToxoDB (r. 48) (96) using the default commands for paired end reads except that the maximum intron length threshold was set at 5000 bases. We checked for mapping quality with Samtools (v.1.7) (102) and visualised subsets of mapped reads via IGV (103). Differential gene expression analysis was performed with featureCounts (v.1.6.2) (104) and limma/voom (v.3.46.0) (42) as previously described (105). We set a significance threshold of 0.05 or less for the adjusted p-value using the Benjamini and Hochberg method, and required a minimum fold change of at least 2 for the gene to be considered differentially expressed. Differential splicing analysis was performed with ASpli (v.1.5.1) (53) using the default commands as previously described (106).

We used the same significance threshold as used for the differential expression analysis. Additionally, the percent intron retention (PIR) or percent spliced in (PSI) metric scores, which are junction inclusion indexes that provide additional evidence for splicing, had to equal or exceed 10% in difference and 2-fold change for the event to be considered differentially spliced. GO enrichment for differential expression was analysed with GSEA (107) using gene ontology terms extracted from ToxoDB (96). GO enrichment for differentially spliced genes was carried out directly in ToxoDB (96) (www.toxodb.org) using the integrated GO tool. In both cases, we required the normalised p-value to be smaller than 0.05 and FDR q-value of less than 0.25 for the term to be considered statistically significant. Transcript productivity was analysed using our in-house tool (gitlab.com/e.mchugh). Proportional Venn diagrams and heatmaps were drawn using BioVenn (108) and pheatmap (109) respectively. All motif analyses were performed with the MEME suite (v.5.3.0) (110) of tools. Motif discovery of AS junctions was done using MEME (55) using the default presets for discriminative mode. For input, we extracted and used the 300 bases coding strand region centred on the 5’ or 3’ splice site of differential (primary) and non-differential (control) IR events. We only considered motifs that were enriched in the primary sequences with an E-value of 0.05 or lower. The positional probability frequency plots of motifs were obtained using CentriMo (110) using the default presets.

## 3.6 Data availability

The RNA-seq data generated and analysed during this study are available from the SRA repository (PRJNA738301).

## 3.7 Supplementary material

**Text S1.** List of PCR primers used in the study.

**Fig S1.** Representative IF images of *Tg*SR-HA-KO bradyzoite parasites.

**Fig S2.** MD plots of gene expression for ablated *Tg*SR and putative *Tg*SR kinase parasites.

**Fig S3.** IGV snapshots of *Tg*SR-HA-KO RNA-seq reads mapping to *Tg*SR genes.

**Fig S4.** Bar graphs of over-represented GO terms in differentially spliced genes.

**Fig S5.** Relationship between differentially spliced and differentially expressed genes.

## Acknowledgements

Microscopy experiments were performed at the Biological Optical Microscopy Platform at the University of Melbourne. The authors thank Aaron Jex (WEHI) for helpful discussions. This work is funded by grants from the Australian National Health and Medical Research Council (Grant 1165354) and the Australian Research Council (DPDP160100389). VVL was funded by a Melbourne Research Scholarship from The University of Melbourne.

---

**Figure S1.**
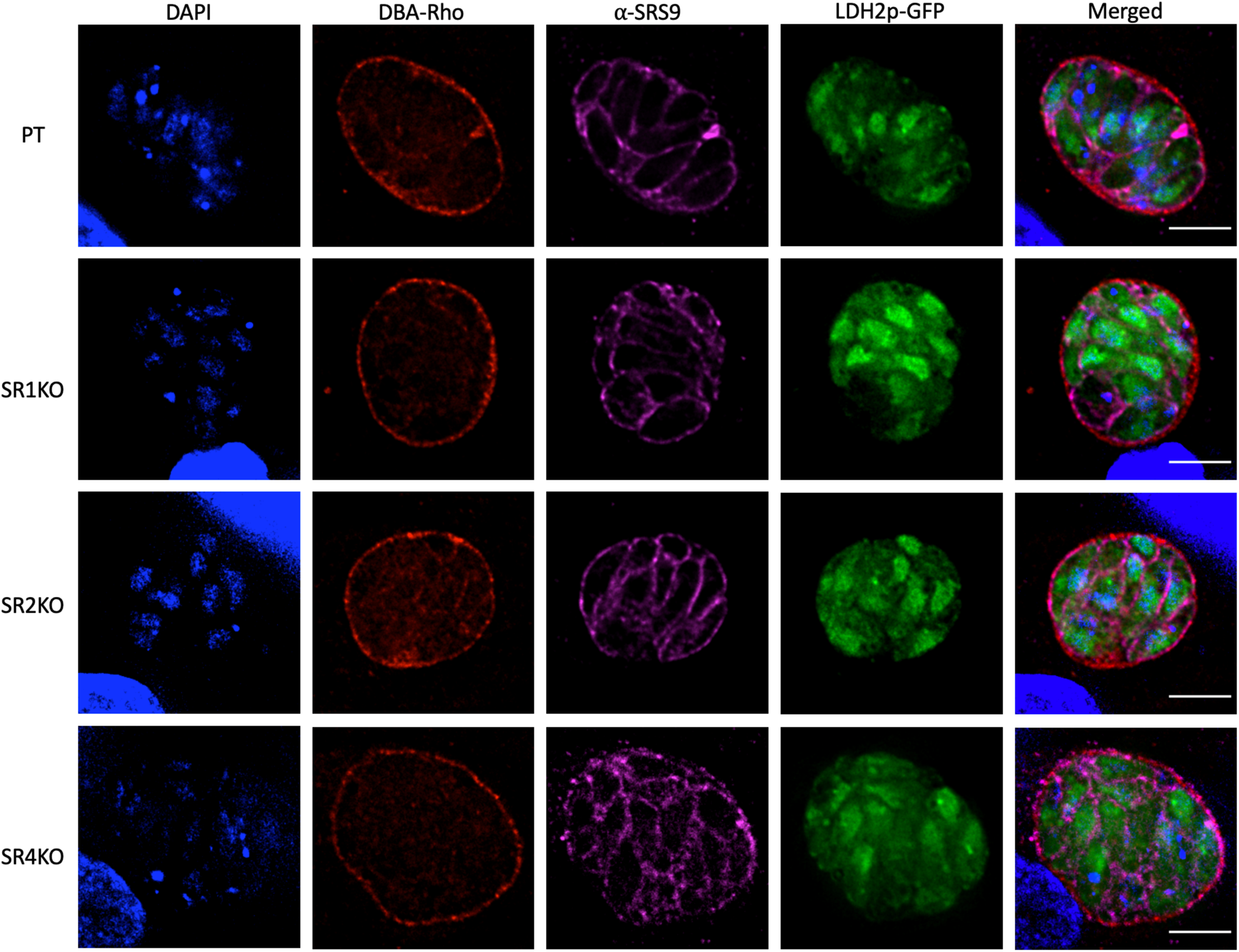
Representative immunofluorescence microscopy images of *Tg*SR-HA-KO bradyzoite parasites stained with DAPI (blue), Rhodamine-Dolichos Biflorus Agglutinin (red) and anti-SRS9 (magenta). Green is GFP which is expressed under the promoter of LDH2. Bradyzoites were differentiated using alkaline induction for 7 days prior to imaging. Images are wide-field deconvoluted of single z stacks. Scale bars = 3 µm.

**Figure S2.**
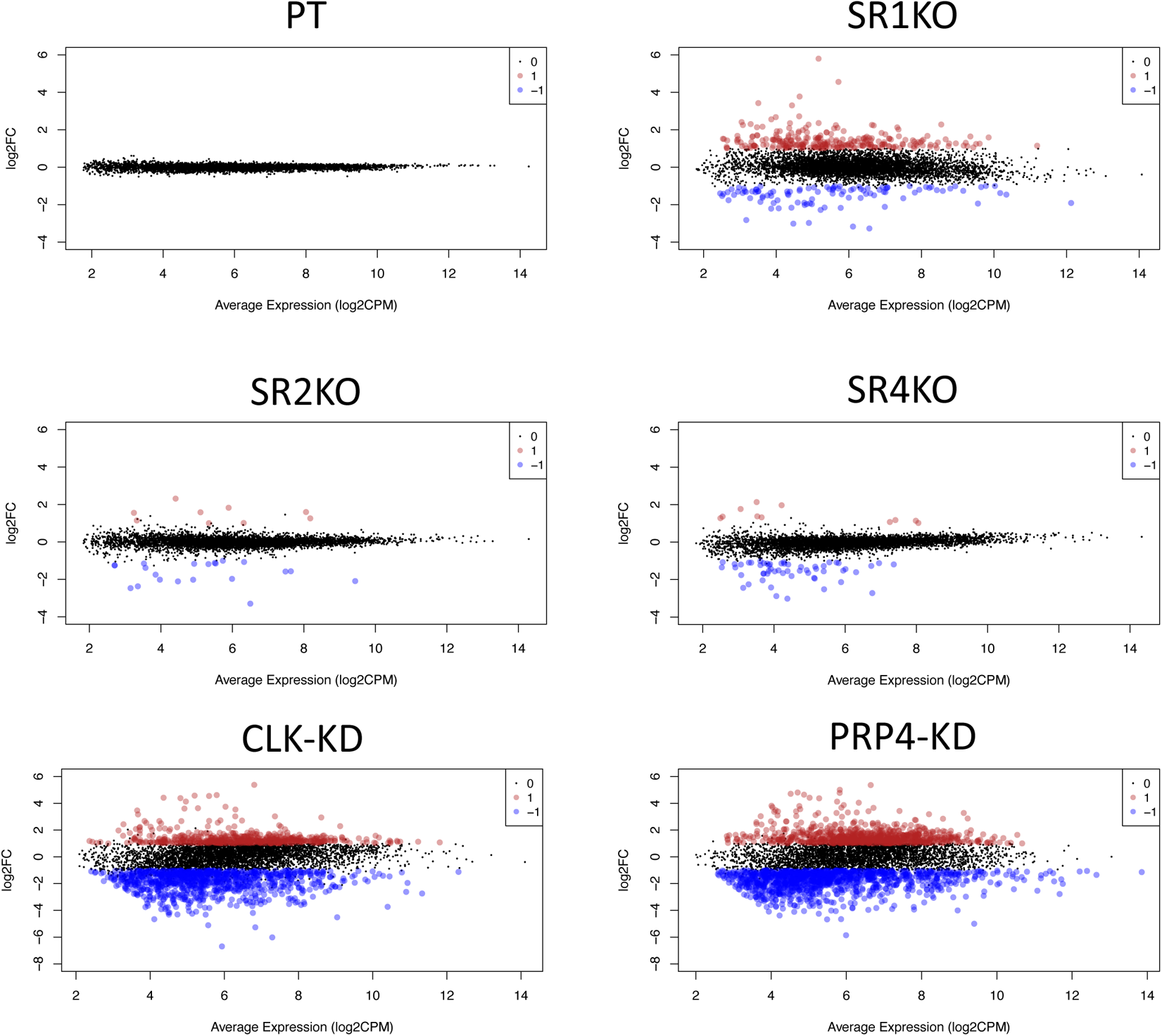
MD plots of gene expression for ablated *Tg*SR and putative *Tg*SR kinase parasites. Differentially expressed genes that were downregulated (−1) or upregulated (1) with an adjusted p-value of ≤ 0.05 and fold change of ≥ 2 are highlighted.

**Figure S3.**
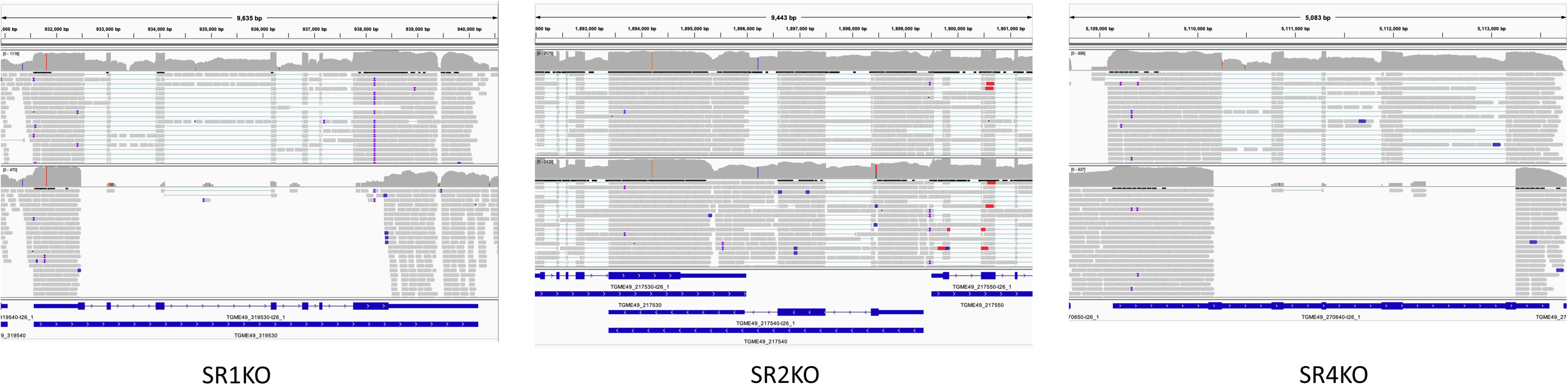
IGV snapshots of RNA-seq reads from parental (top) and each *Tg*SR-HA-KO (bottom) parasite lines mapping to their respective *Tg*SR genes. *Tg*SR1 and *Tg*SR4 KO mutants show no reads while *Tg*SR2 show an altered mapping at the site of cassette insertion near the 5’ end of the gene.

**Figure S4.**
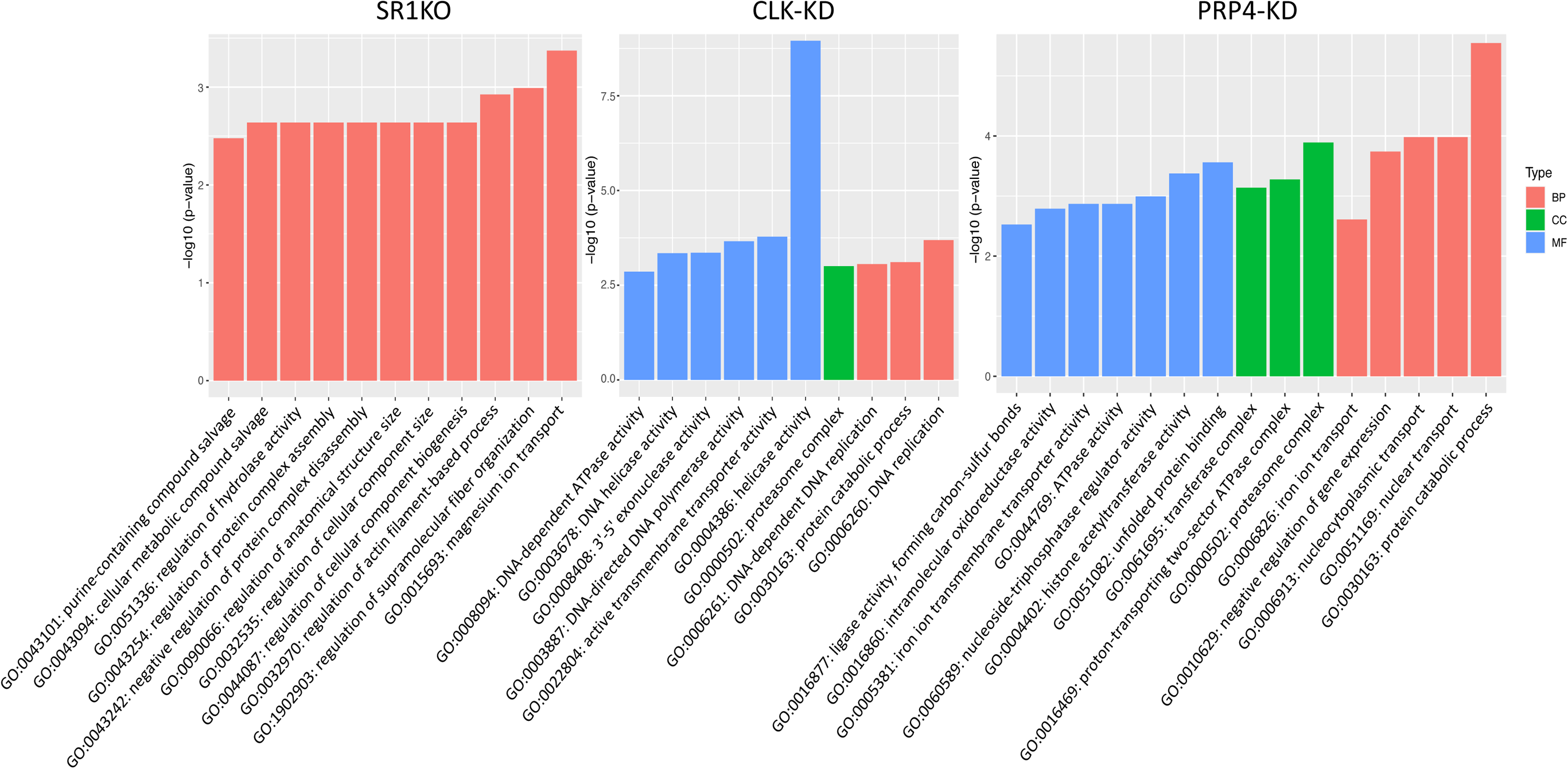
Bar graphs of GO terms that were over-represented in genes that were differentially spliced for each parasite genotype/condition, where present. Only terms with a p-value of ≤ 0.05 and FDR of ≤ 0.25 were considered over-represented.

**Figure S5.**
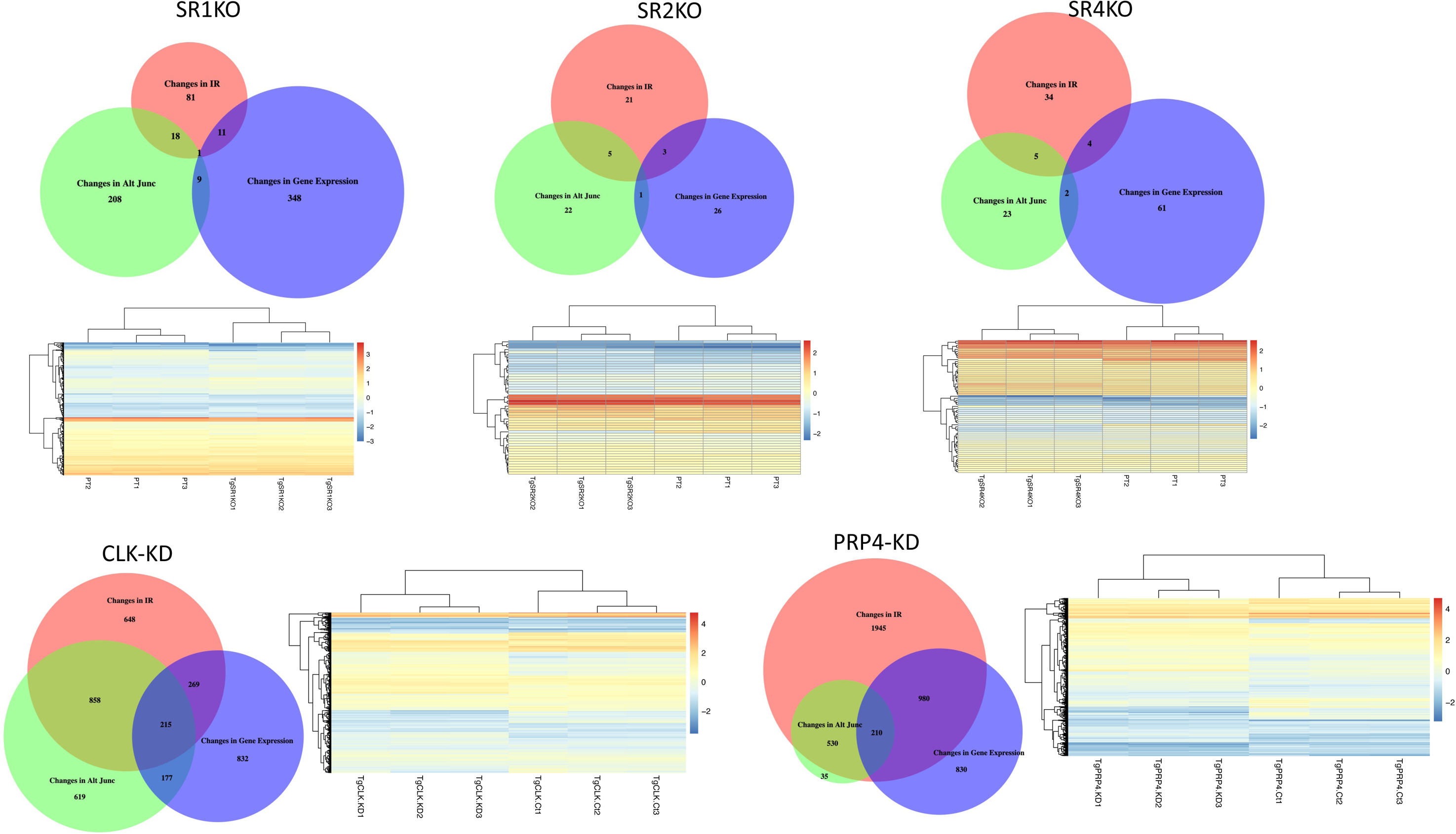
Relationship between differentially spliced and differentially expressed genes in ablated *Tg*SR and putative *Tg*SR kinase parasites. The proportional Venn diagrams show the overlap between differentially spliced and differentially expressed genes. Differentially spliced genes are separated based on whether there are changes to intron retention (IR) or other types of alternative splicing (AS). Accompanying heatmaps show the changes in normalised expression gene expression for differentially spliced genes in each sample.

